# A random mutagenesis screen enriched for missense mutations in bacterial effector proteins

**DOI:** 10.1101/2024.03.14.585084

**Authors:** Malene L. Urbanus, Thomas M. Zheng, Anna N. Khusnutdinova, Doreen Banh, Harley O’Connor Mount, Alind Gupta, Peter J. Stoigos, Alexei Savchenko, Ralph R. Isberg, Alexander F. Yakunin, Alexander W. Ensminger

## Abstract

To remodel their hosts and escape immune defenses, many pathogens rely on large arsenals of proteins (effectors) that are delivered to the host cell using dedicated translocation machinery. Effectors hold significant insight into the biology of both the pathogens that encode for them and the host pathways that they manipulate. One of the most powerful systems biology tools for studying effectors is the model organism, *Saccharomyces cerevisiae*. For many pathogens, the heterologous expression of effectors in yeast is growth inhibitory at a frequency much higher than housekeeping genes, an observation ascribed to targeting conserved eukaryotic proteins. Abrogation of yeast growth inhibition has been used to identify bacterial suppressors of effector activity, host targets, and functional residues and domains within effector proteins. We present here a yeast-based method for enriching for informative, in-frame, missense mutations in a pool of random effector mutants. We benchmark this approach against three effectors from *Legionella pneumophila*, an intracellular bacterial pathogen that injects a staggering >330 effectors into the host cell. For each protein, we show how *in silico* protein modeling (AlphaFold2) and missense- directed mutagenesis can be combined to reveal important structural features within effectors. We identify known active site residues within the metalloprotease RavK, highly conserved residues in SdbB, and previously unidentified functional motifs within the C-terminal domain of SdbA. We show that this domain has structural similarity with glycosyltransferases and exhibits *in vitro* activity consistent with this predicted function.

## Introduction

For many bacterial pathogens, host manipulation derives from the collective activity of large numbers of translocated proteins (“effectors”) that are injected into the host cell using dedicated secretion machinery. A striking example is the gram-negative bacterium, *Legionella pneumophila*, the causative agent of Legionnaires’ disease, an often-fatal severe pneumonia that naturally replicates in freshwater protozoa. *L. pneumophila* encodes for the largest effector arsenal described to date (>330 effectors per isolate, or roughly 10% of the proteome) (Burstein *et al*. 2009; Huang *et al*. 2011; Zhu *et al*. 2011), which is injected into the host cell using the Dot/Icm type IVB secretion system (Segal *et al*. 1998; Vogel *et al*. 1998). *L. pneumophila* effectors modulate host vesicle trafficking, post-translational modification, protein translation, autophagy, vacuolar function, and the cytoskeleton to avoid lysosomal fusion and to establish a replicative, neutral pH vacuole (Isberg *et al*. 2009; Escoll *et al*. 2013; Sherwood and Roy 2016; Qiu and Luo 2017). While over 50 effectors have been studied, the majority of these remain uncharacterized (Finsel and Hilbi 2015).

Determining the function of the >330 effectors and the role they play in establishing the *Legionella*-containing vacuole is complicated by extensive genetic redundancy within the effector arsenal (O’Connor *et al*. 2011) and the lack of predicted conserved domains or functions for many substrates (Gomez-Valero *et al*. 2011, 2014; Burstein *et al*. 2016). In a study comparing genomes from 38 *Legionella* species, only half of the predicted effectors contained conserved domains and many of these are of uncharacterized function (Burstein *et al*. 2016).

Although the amino acid sequence of many effectors may not yield obvious clues to their function, some effectors have structural homology to characterized proteins or domains, along with conserved active site motifs or other signature motifs (Toulabi *et al*. 2013; Morar *et al*. 2015; Wong *et al*. 2015; Urbanus *et al*. 2016; Pinotsis and Waksman 2017; Kozlov *et al*. 2018). Looking beyond *L. pneumophila*, over 18,000 effector genes have been predicted across the entire *Legionella* genus (Burstein *et al*. 2016; Gomez-Valero *et al*. 2019). This number suggests both a wealth of novel effector activities and host biology waiting to be discovered.

We set out to develop a method to efficiently identify important motifs or amino acid residues in uncharacterized *Legionella* effectors by random mutagenesis and selection for loss-of-function mutations to facilitate the prediction of mechanism and function. Approximately 10% of the effectors severely inhibit yeast growth when overexpressed (Campodonico *et al*. 2005; Felipe *et al*. 2008; Heidtman *et al*. 2009; Shen *et al*. 2009; Guo *et al*. 2014; Urbanus *et al*. 2016), such that loss-of-function by random mutagenesis can be selected for as an alleviation of the yeast growth defect. However, a random mutant pool contains a multitude of mutations that can potentially cause a loss-of-function phenotype, such as frameshift, nonsense and missense mutations in the effector or regulatory elements such as the promoter region. Most of these mutations are uninformative as only missense loss-of-function mutations in the effector are relevant. To enrich for full-length missense clones, we used a C-terminal in-frame His3 fusion and selected for histidine prototrophic yeast requiring the presence of a full-length protein. A similar strategy (C-terminal His3 fusions) was previously shown to enrich for in-frame human open reading frames amongst a randomly primed pool of cDNAs cloned into a yeast expression vector (Holz *et al*. 2001). Here, we benchmark this in-frame mutagenesis approach against three *L. pneumophila* effectors previously shown to inhibit yeast growth: SdbA and SdbB, whose functions remain uncharacterized and RavK, a previously described metalloprotease. We show our approach identifies active site residues within RavK (Liu *et al*. 2017) highly conserved residues in SdbB, and previously unidentified functional motifs in the C-terminal domain of SdbA. These motifs are part of the donor and acceptor binding regions of glycosyltransferases, which we show the C-terminal domain of SdbA shares homology with. Finally, we show that a C-terminal fragment exhibits *in vitro* activity consistent with this predicted function.

## Results

### A random mutagenesis screen to identify regions important for bacterial effector function

To efficiently screen for randomly generated mutations in *Legionella* effectors that cause a loss-of-function phenotype and represent full-length protein rather than frameshift or nonsense mutations, we applied the method of Holz and colleagues to select for expression of full-length proteins in yeast (Holz *et al*. 2001). We cloned the yeast *HIS3* gene in frame behind yeast- cytotoxic *Legionella* effectors on a high-copy galactose-inducible yeast expression plasmid.

After confirming that the His3 fusion does not interfere with the yeast-cytotoxic phenotype, and therefore likely does not interfere with effector function, we generated a pool of random mutants using the *E. coli* mutator strain XL1-Red. We then transformed this mutant plasmid pool to yeast and monitored growth on different media types to assess the number of transformable plasmids (- ura/glucose) and the efficiency of the random mutagenesis step (-ura/galactose) selecting all mutations that caused a loss of function (Fig 1). To specifically select for full-length missense loss-of-function mutants, we grew the transformed pool on medium with galactose and lacking uracil and histidine (-ura-his/galactose), which requires the production of full-length effector- His3 fusion protein and mutations in the effector that disrupt its activity.

**Figure 1:**
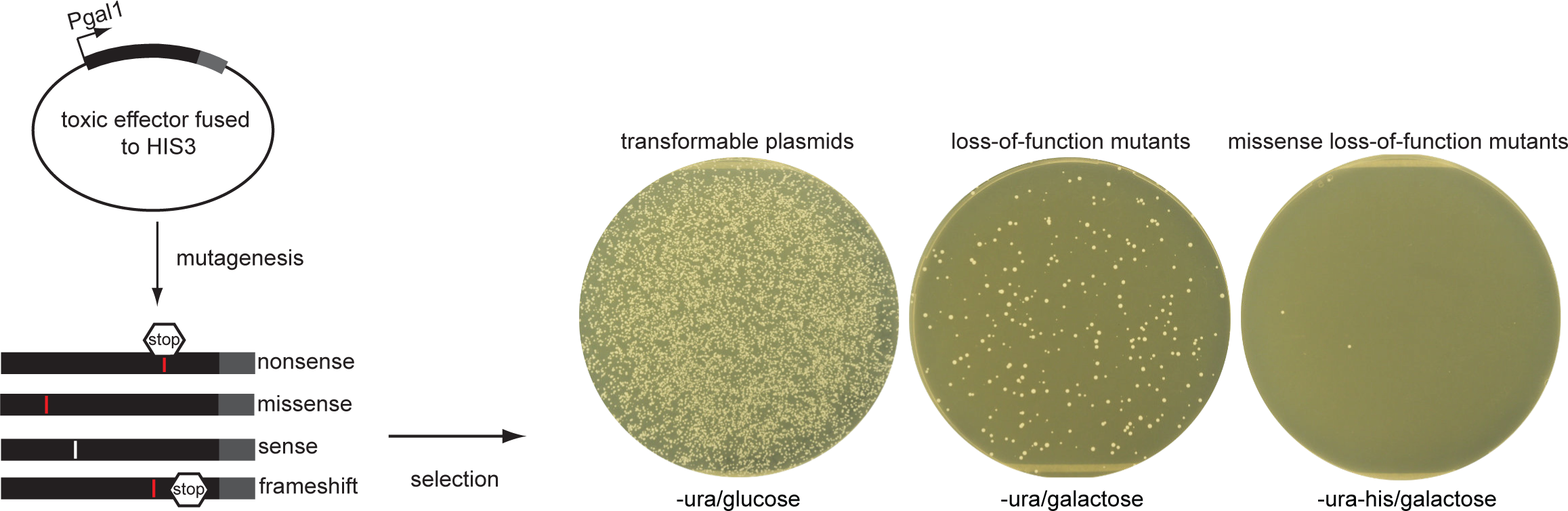
**A method to enrich full-length missense mutants in a random mutagenesis screen**. *Legionella* effectors with a severe growth phenotype in yeast are fused in frame with the yeast *HIS3* gene in a high-copy yeast expression vector with galactose-inducible expression and an uracil selection marker. The plasmid is randomly mutated in a *E. coli* mutator strain, such as XL1-red. The resulting mutant pool contains a variety of mutants; sense, nonsense, missense and frameshifts, of which the latter three can cause loss-of-function phenotypes. The plasmid pool is transformed to *S. cerevisiae* strain BY4742 and grown under conditions that select all transformable plasmids (–ura/glucose), all loss-of-function mutations (-ura with galactose induction) and only full-length proteins containing missense loss-of-function mutations (-ura-his with galactose induction).

### Missense loss-of-function screen identifies conserved SdbB amino acid residues

As a proof of principle, we looked at SdbB which severely inhibits yeast growth (Fig 2A) (Heidtman *et al*. 2009) and is part of the SidB paralog family, whose members are predicted to be lipases from the α/β hydrolase enzyme family (Luo and Isberg 2004). After verifying that the SdbB-His3 fusion was still capable of inhibiting yeast growth (Fig 2A) we created a random mutagenesis pool of SdbB-His3 and quantified the number of colony-forming units (CFUs) on the different selection media. While 1.89% of the transformable plasmids carried a mutation that allowed for growth on SD-ura/galactose medium indicating some type of loss-of-function mutation (Fig 2B), only 0.03% of the transformable plasmids carried a mutation allowing growth on SD-ura/-his/galactose medium which requires the expression of full-length fusion protein.

**Figure 2:**
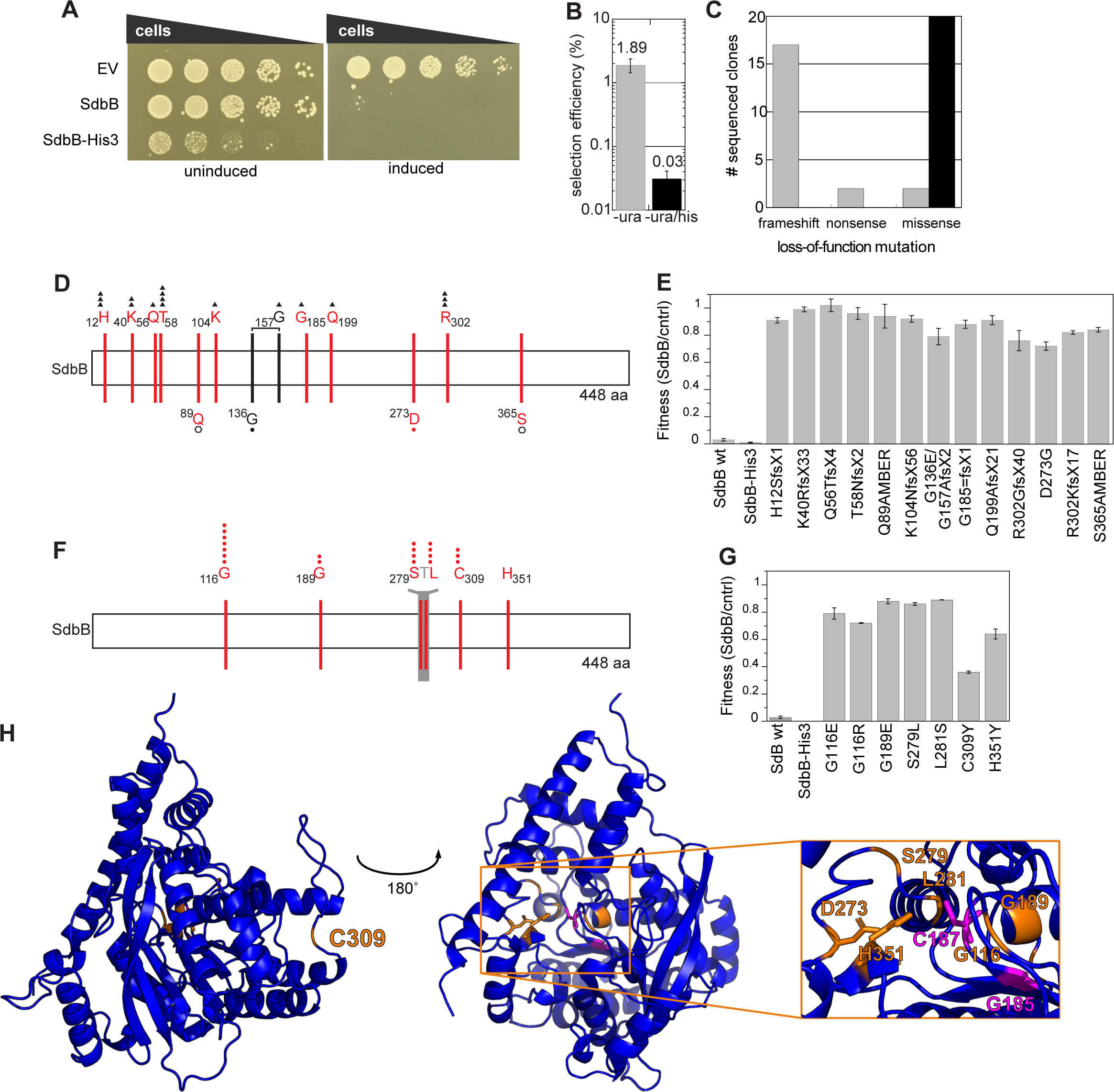
The full-length ORF enriched random mutagenesis screen identifies missense mutations in the putative active site pocket. A. SdbB (lpg2482) and SdbB in-frame fusion with His3 show a severe growth defect in a yeast spot dilution assay. **B** The percentage of loss-of- function mutation in the pool of transformable plasmids selected on medium lacking uracil (-ura, grey) and lacking uracil and histidine (-ura-his, black) with galactose induction. The average and standard deviation of three independent replicates are shown. **C** Occurrence of frameshift, nonsense and missense mutations in 20 sequenced clones selected on -ura and -ura-his with galactose induction. **D** A schematic representation of loss-of-function clones selected on –ura with galactose induction. Mutations recovered from the same clone are shown in black, connected by a line while single mutations are shown in red. Mutation type is indicated as a closed triangle (frameshift), open hexagon (nonsense) or closed circle (missense), and the number of symbols reflects the occurrence of the mutation in the dataset. **E** The fitness of wild- type SdbB, His3 fusion and loss-of-function mutations (selected on –ura/galactose) compared to empty vector controls in liquid growth assays confirms the loss-of-function phenotype for the SdbB random mutagenesis clones. The average and standard deviation of three technical replicates are shown. **F** A schematic representation of SdbB loss-of-function clones selected on - ura-his/galactose. Missense loss-of-function mutations are shown in red with a closed circle, the number of symbols reflects the occurrence of the mutation in the dataset. Amino acid residues shown for presentation purposes are shown in grey. **G** The fitness of wild-type SdbB, His3 fusion and loss-of-function mutations (selected on -ura,-his/galactose) compared to empty vector controls in liquid growth assays confirms the loss-of-function phenotype for the SdbB random mutagenesis clones. The average and standard deviation of three technical replicates are shown. **H** The missense mutations identified by the random mutagenesis screen (D,F) are shown in orange on an Alphafold2 model of SdbB (AF-Q5ZSN5-F1-model_v4), and residues from the putative active site 185-GxCxG189 not captured by the screen are shown in magenta. Putative catalytic triad C187-D273-H351 residies are shown with sticks and the box shows the enlargement of the putative catalytic site. The AlphaFold2 model was visualized using Pymol 2.2 (Schrödinger).

The efficiency of the histidine selection step was verified by sequencing 20 clones from each condition. The loss-of-function clones selected on –ura/galactose consisted of 16 frameshift mutations, 2 nonsense mutations, 1 combination of a missense and frameshift mutation and 1 missense mutation in the *sdbB* gene (Fig 2C). In contrast, the loss-of-function clones selected on –ura/-his/galactose only contained missense mutations (Fig 2D,F). For both conditions a number of mutations were recovered several times suggesting that the mutational screen is reaching saturation. To confirm that the identified mutations indeed rescued SdbB toxicity in yeast, we compared the fitness of mutants to an empty vector control in a liquid growth curve assay (see Experimental methods). The wild-type and His3-fused SdbB almost completely inhibited yeast growth, while the SdbB mutants showed a fitness of 60-90% compared to an empty vector control (Fig 2E and 2G).

The positions of the frameshift and nonsense mutations in SdbB (Fig 2D) indicate that a large part of the protein is required for function, as even a nonsense mutation at S365, 84 amino acid residues from the C-terminus, almost completely rescued activity. The missense loss-of-function mutations (Fig 2D and 2F) target four amino acid residues (G116, G189, D273 and H351) that are invariant in SdbB orthologs from *Legionella pneumophila* and other *Legionella* species (Burstein *et al*. 2016) (Fig S1), suggesting they are essential for function or structure.

The G198E mutation is part of the GXS/CXG motif that is invariant across the SdbB orthologs and is predicted to align with the so-called nucleophile elbow of the nucleophile-acid-base triad of the α/β hydrolase active site (Brenner 1988; Ollis *et al*. 1992; Schrag and Cygler 1997; Marchler-Bauer *et al*. 2017) (Fig S2). When the mutations are mapped onto the SdbB AlphaFold2 model (Jumper *et al*. 2021; Varadi *et al*. 2021) the majority localize in the pocket with the putative catalytic cysteine (C187), including the invariant D273 and H351 residues captured in the screen suggesting they are the remaining residues of the catalytic triad (Fig 2H).

Thus, the SdbB example demonstrates that functionally important amino acid residues can be identified using the random mutagenesis method in conjunction with the histidine selection for full-length protein. Importantly, this approach significantly reduces the number of sequenced clones required to identify amino acid residues or regions of interest, by approximately 60-fold in the case of SdbB.

### Missense loss-of-function screen identifies the active site of the characterized effector RavK

To benchmark the missense loss-of-function screen on a characterized effector, we looked at RavK which also severely inhibits yeast growth (Heidtman *et al*. 2009). RavK was recently shown to be a small, soluble metalloprotease that specifically cleaves host actin, and mutation of the active site motif HExxH abolishes all activity and yeast toxicity (Liu *et al*. 2017). The RavK- His3 fusion was still able to inhibit yeast growth (Fig 3B) and subjected to random mutagenesis and selection for loss-of-function mutants on medium lacking histidine. Of the 14 loss-of-function clones we sequenced, one clone contained a large, in-frame deletion from amino acid residue 70 to residue 166, which was unexpected but confirms the strength of the histidine selection for full-length protein. All other loss-of-function clones were caused by single point mutations resulting in missense mutations (Fig 3A). In the fitness assay, wild-type and His3- fused RavK almost completely inhibited yeast growth, while the RavK mutants displayed a fitness of 70-90% compared to the empty vector control. The loss-of-function mutations all map to the first half of RavK, suggesting that the N-terminal half of RavK is essential for RavK function. This is in agreement with Liu and colleagues (Liu *et al*. 2017), who identified the active site motif (_95_HExxH_99_) in the N-terminal half of the protein and found that the 50 C-terminal residues of RavK can be deleted without any effect on its activity on actin.

**Figure 3:**
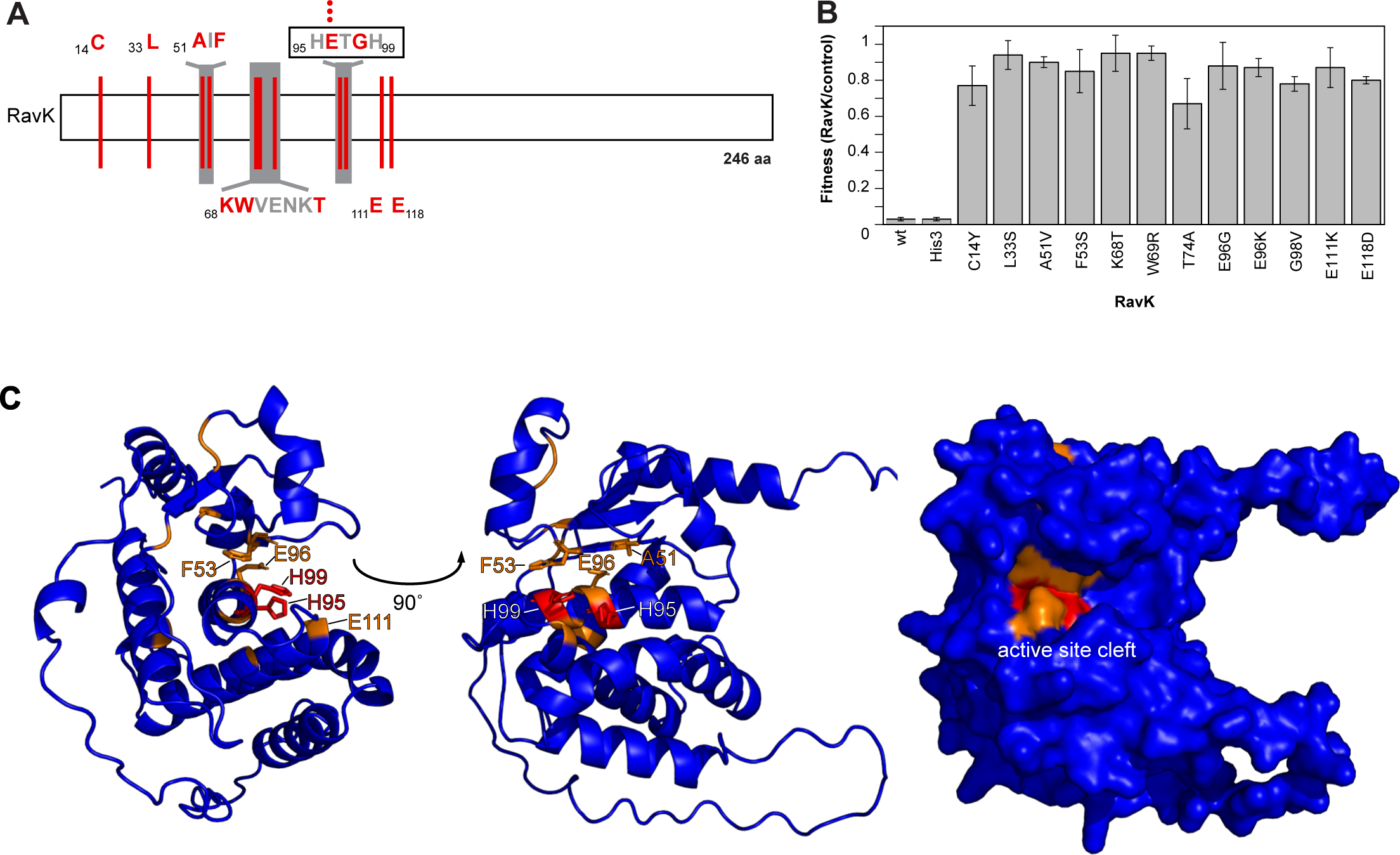
RavK random mutagenesis captures residues lining the active site cleft. A. A schematic representation of the mutations causing a RavK loss-of-function phenotype when expressed in yeast. Mutated residues are shown in red, amino acid residues shown for presentation purposes are in grey and the number of symbols reflects the occurrence of the mutation in the dataset. **B** The growth fitness of wild-type RavK, RavK-His3 and loss-of-function mutants normalized to empty vector controls (see Methods) confirms the loss-of- function phenotype for the RavK random mutagenesis clones. The average and standard deviation of three technical replicates are shown. **C** The Alphafold2 model of RavK (AF- Q5ZWW5-F1-model_v4.pdb) with missense mutations shown in orange. The histidine residues of the active site motive _95_HExxH_99_ (Liu *et al*. 2017) not captured by the screen are shown in red. The residues in the active site cleft are shown as sticks. The AlphaFold2 model was visualized using Pymol 2.2 (Schrödinger).

Four of the loss-of-function mutations targeted the active site motif H_95_ExxH_99_, three of which are mutations to E_96_. Our screen also identified several residues outside of this motif that are critical for RavK function. To investigate why these might be functionally important, we performed an HHpred analysis which looks for structural homologs of proteins (Zimmermann *et al*. 2018). HHpred identified many hits with homology to the HExxH metalloprotease motif.

Among the top five HHpred hits are three small soluble metalloproteases or minigluzincins, anthrax lethal factor and a zinc dependent peptidase from the M48 family (Dalkas *et al*. 2010; López-Pelegrín *et al*. 2013; LópezLJPelegrín *et al*. 2014) (Fig S3). Notably, some of the other loss-of-function mutations occur in areas that have homology with structural elements in the minigluzincins that contribute to the active site cleft (López-Pelegrín *et al*. 2013) (Fig S3).

Indeed, when the missense mutations are mapped on the RavK AlphaFold2 model (Jumper *et al*. 2021; Varadi *et al*. 2021) (Fig 3C) they cluster around the active site HExxH including the top rim of the active site cleft.

### The C-terminal domain of SdbA is a putative glycosyltransferase

SdbA is a member of the SidB paralog family (Luo and Isberg 2004). While the function of SdbA remains undefined, experimental evolution of *Legionella* in mouse macrophages selected for parallel SdbA nonsense and frameshift mutations in three out of four independent lineages (Ensminger *et al*. 2012), suggesting that its normal function partially restricts growth in this accidental host. While the N-terminal domain of SdbA has homology with SidB (Luo and Isberg 2004), the additional C-terminal domain does not have significant sequence homology to other known proteins (data not shown). SdbA completely inhibits yeast growth when heterologously expressed (Heidtman *et al*. 2009), making the missense loss-of-function screen an informative tool by which to identify functional residues that might suggest a specific activity inside the eukaryotic cell.

The missense loss-of-function screen identified 19 mutations in 24 sequenced clones targeting 17 amino acid residues in SdbA (Fig 4A). In contrast to the smaller proteins SdbB and RavK, the SdbA results included several double mutants. In some cases, it is not clear how much each mutation contributes to the phenotype, considering that the fitness of the double mutants is not dramatically different from the single loss-of-function mutations (Fig 4B). However, all the single mutations that lead to a loss-of-function phenotype fall in the C-terminal domain and mainly concentrate in two regions: _541_GGTGHIS_547_ and _957_GGLSVME_963_. An HHpred homology search (Zimmermann *et al*. 2018) predicted with high confidence that the C-terminal domain is a glycosyltransferase of the GT-B fold. When comparing the SdbA C-terminal domain with the sequence of *E. coli* MurG, a well-studied member of the GT-B fold glycosyltransferase family, the two mutated regions align with the G-loop 1 and a consensus region in GT-B fold superfamily involved in binding the donor molecule (Ha *et al*. 2000; Hu *et al*. 2003; Crouvoisier *et al*. 2007) (Fig 4C). Glycosyltranferases hydrolyze UDP-sugar donor molecules and transfer the sugar to the acceptor molecule – which can be a variety of molecules such as small molecules, lipid or proteins (Lairson *et al*. 2008). In MurG amino acid residues A263, L264, L265, E268, Q287 and Q288 contact the donor molecule UDP-GlcNAc (Hu *et al*. 2003) (Fig4C) while the G-loop 1 is thought to be involved in acceptor molecule binding (Ha *et al*. 2000). Notably, mutations in these motifs abrogate MurG enzymatic activity, including mutation of the residues H18 and E268 (Hu *et al*. 2003; Crouvoisier *et al*. 2007), whose corresponding residues in SdbA (H545 and E963) were found to be mutated in our screen. Fig 4D shows the single missense mutants abrogating SdbA activity mapped onto the AlphaFold2 model (Jumper *et al*. 2021; Varadi *et al*. 2021)of a C-terminal fragment (amino acid residues 510-1050) with the residues H545 and E963 highlighted in yellow (43) To test whether the C-terminal domain of SdbA is indeed a glycosyltransferase, we purified the C-terminal fragment 510-1050 and the equivalent of the MurG E268A inactive mutant in SdbA (E963A) and tested several UDP-sugars as substrate using the UDP-Glo assay (Fig 4E). Glycosyltransferases can hydrolyze UDP-sugars in the absence of an acceptor molecule (with water acting as an acceptor in the reaction)(Sheikh *et al*. 2017; Vicente *et al*. 2023). Indeed wild-type SdbA 510-1050 hydrolyzes the UDP-GlcNAc donor while the E963A mutant cannot (Fig 4E), confirming that SdbA is a glycosyltransferase with specificity for UDP-GlcNAc and that a mutation in the E963 residue identified by the missense loss-of-function screen abrogates that activity. Using the same assay, we determined the kinetic parameters of UDP-GlcNAc hydrolysis by SdbA (Fig 4F), which revealed high affinity (low micromolar Km) of this enzyme to UDP-GlcNAc suggesting efficient glycosylation activity *in vivo*.

**Figure 4:**
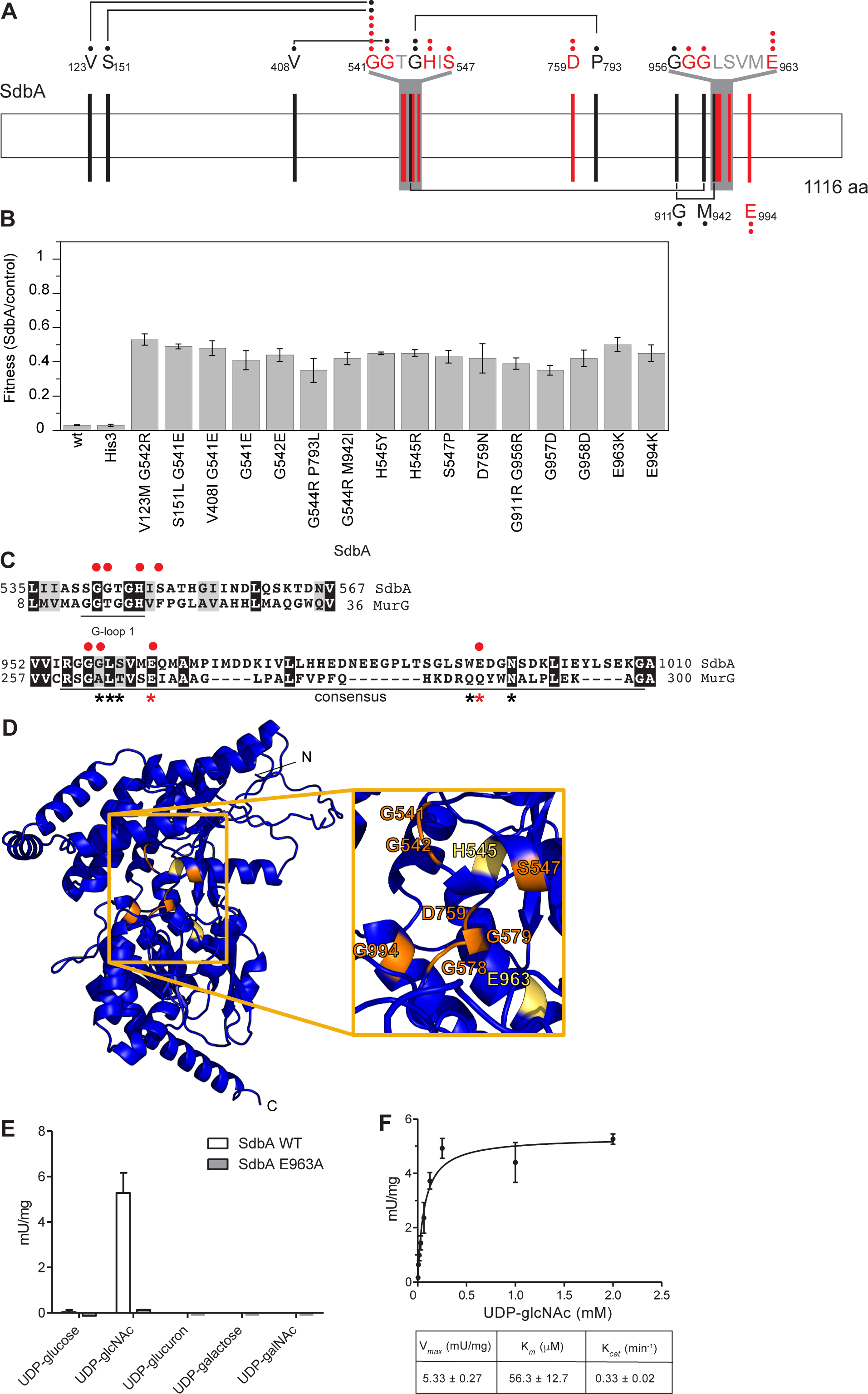
The C-terminal domain of SdbA is a putative glycosyltransferase domain. **A** A schematic representation of the mutations causing a SdbA loss-of-function phenotype when expressed in yeast. Mutations recovered from the same clone are shown in black, connected by a line and single mutations alleviating a SdbA induced growth defect are shown in red. Amino acid residues shown for presentation purposes are shown in grey. Black and red closed circles indicate the number of times the mutation was identified. **B** The growth fitness of wild-type SdbA and loss-of-function mutants normalized to empty vector controls (see Methods for details). SdbA and SdbA-His3 display a severe growth defect, while the loss-of-function mutations rescue growth up to 50% of the empty vector control. The average and standard deviation of three technical replicates are shown. **C** HHpred alignment of SdbA with 1F0K (Ha *et al*. 2000) (*E. coli* MurG). The G-loop 1 and consensus sequence are shown with identical residues (black) and similar residues (grey) highlighted. SdbA amino acid residues that when mutated abrogate activity are indicated by a red closed circle. Amino acid residues in the MurG consensus sequence contacting UDP-GlcNac are indicated by a black star and a red start if mutations abrogate MurG activity. **D** The Alphafold2 model of SdbA (AF-Q5ZYT6-F1- model_v4.pdb) with missense mutations shown in orange. Residues H545 and E963, corresponding to residues H18 and E268 in MurG, are shown in yellow (43). The AlphaFold2 model was visualized using Pymol 2.2 (Schrödinger). **E** The SdbA glycosyltransferase domain uses GlcNac as a donor substrate. Donor substrate specificity was tested by incubating 0.09 µM of purified fragment (residues 510-1050) of wild-type SdbA or an inactive mutant (E963A) with 100 µM UDP-glucose, UDP-glcNAc, UDP-glucuron, UDP-galactose or UDP-galNAc for 1h at 30°C. The hydrolase activity of the UDP-substrate was detected as the release of UDP by the UDP-Glo assay (Promega) after 20 min of incubation with UDP-Glo detection reagent. Luminescence was measured using a SpectraMax M2 platereader. Three technical repeats were performed per reaction. **F** Determination of the kinetic parameters for GlcNac hydrolysis by SdbA. Reactions with 0.16 µg purified wild-type C-terminal domain of SdbA (residues 510- 1050) and a range of 0.0039 – 2 mM GlcNac and were incubated for 1 h at 30°C. GlcNac hydrolysis was measured using the UDP-Glo glycosyltransferase assay as described above; three technical replicates were performed per reaction. Kinetic parameters were determined by non- linear curve fitting from the Michaelis-Menten plot.

Taken together, the missense loss-of-function screen identified critical residues in the C-terminal domain of SdbA, which together with HHpred analysis predict that SdbA contains a glycosyltransferase domain. The discovery and confirmation of the C-terminal glycosyltransferase domain allows for targeted follow-up experiments to elucidate the action and target of SdbA on eukaryotic host cells.

## Discussion

Our observations demonstrate that the combination of a random mutagenesis loss-of-function screen with a selection for full-length protein is very effective in specifically selecting loss-of-function missense clones. In fact, all but one of the clones recovered in our assay contained missense mutations, while the remaining one contained an in-frame deletion which included the active site of RavK. The screens correctly identified the previously described active site of RavK, and the predicted active site nucleophile motif and the remaining residues of the catalytic triad of SdbB. Surprisingly, the N-terminal α/β hydrolase domain of SdbA did not accumulate loss-of-function mutations, but the C-terminal domain appears to be crucial for the function of SdbA. The prediction that the C-terminal domain of SdbA is a glycosyltransferase was supported by *in vitro* activity towards UDP-GlcNAc and provides a way forward to study this *Legionella* effector.

To effectively apply the random mutagenesis missense enrichment selection or to extend this approach to bacteria and mammalian cells, a number of considerations need to be taken into account. First, the protein of interest must have a selectable loss-of-function phenotype such as alleviation of growth inhibition. Growth fitness is a universal phenotype that is easy to measure in bacteria, yeasts and mammalian cell lines. Second, the C-terminal fusion of a selection marker must not interfere with protein function. If the function of protein of interest is inhibited by the C-terminal fusion, it could potentially be overcome by introducing linker regions of varying length and flexibility (Chen *et al*. 2013) or by using a cleavable linker such as the ubiquitin K0 mutant that is processed by cytosolic deubiquitinases in eukaryotic cells (Bachran *et al*. 2013).

Similarly, the C-terminal selection marker must be able to function as a fusion protein or be liberated by an *in vivo* cleavable linker.

To extend the random mutagenesis missense enrichment selection to bacteria, the chloramphenicol acetyltransferase (CAT) gene which confers resistance to chloramphenicol could be a good candidate as a C-terminal fusion partner. CAT has been successfully used in protein fusions where it conferred chloramphenicol resistance during colony selection as a C-terminal fusion partner, with increased selection efficiency when mutant fusion proteins are soluble (Maxwell *et al*. 1999). In mammalian cell lines, a positive selection marker such as blasticidin S deamidase could be used as a C-terminal fusion partner. Blasticidin S deamidase is functional as a C-terminally fused protein (Suarez and McElwain 2009) and confers resistance against blasticidin, which rapidly inhibits mammalian cell growth at a low dose (Kimura *et al*. 1994). An alternative, if no positive selection marker is available, is GFP, which has been used extensively as a fused localization marker for various cellular compartments and organisms (Margolin 2000; Roessel and Brand 2002; Huh *et al*. 2003). After a standard number of generation doublings, GFP-positive cells, indicative of the presence of full-length protein, can be selected by flow cytometry and cell sorting or by screening for GFP positive colonies in bacteria and yeast. However, these approaches would reduce the screening throughput and/or require specialized equipment.

Our initial results suggest that missense-directed mutagenesis will be a useful tool to help identify potential functions for other bacterial effector proteins, many of which have low sequence homology to characterized proteins (Gomez-Valero *et al*. 2011; Burstein *et al*. 2016). Rather than being replaced by *in silico* protein modeling, we show how the two methodologies complement one another and can be used to identify structural features essential for activity against the eukaryotic cell. In *Legionella pneumophila* alone 10% of the translocated effectors (Campodonico *et al*. 2005; Felipe *et al*. 2008; Heidtman *et al*. 2009; Shen *et al*. 2009; Guo *et al*. 2014; Urbanus *et al*. 2016), severely inhibit yeast growth and are possible candidates for the random mutagenesis missense enrichment screen. Identifying functional residues within uncharacterized effectors is a logical first step towards validating *in silico* protein models, predicting effector activity, and designing protein-protein interaction studies.

## Experimental methods

### In frame effector-His3 fusion by yeast recombinational cloning

The *S. cerevisiae* BY4742 (*MATa, his3*Δ*1, leu2*Δ*0, met15*Δ*0, ura3*Δ*0*; (Brachmann *et al*. 1998)) strains overexpressing *lpg0275*, *lpg0969* and *lpg2482* (*sdbA, ravK,* and *sdbB* respectively) from the high-copy vector pYES2 NT/A (Life Technologies, GAL 1 promoter, N-terminal 6X HIS/Xpress tag and URA3 selectable marker) (Heidtman *et al*. 2009) were used to create the effector-His3 fusion mutants by yeast recombinational cloning. The *S. cerevisiae HIS3* gene was PCR amplified from pAG423GAL-ccdB (Alberti *et al*. 2007) using an effector specific forward primer containing the last 50-60 nucleotides of the effector (minus the stop codon) followed by the first 20-30 nucleotides of the *HIS3* sequence and the pYES-HIS3 reverse primer (Table S1). The resulting PCR products were transformed together with XbaI/PmeI digested pYES2 NT/A lpg0275, lpg0969 or lpg2482 to BY4742 using the high-efficiency LiOAC/PEG method (Gietz and Schiestl 2007) and plated onto SD-uracil with 2% glucose. The resulting transformants were screened by PCR and sequence verified. To confirm that the His3 fusion does not interfere with effector function, the ability of the effector-HIS3 fusion to inhibit yeast growth was tested by comparing the growth BY4742 with empty vector control, the wild-type effector and the effector-HIS3 fusion in a yeast spot dilution assay as described previously (Urbanus *et al*. 2016).

### Selection of loss-of-function mutations

The effector-HIS3 fusion vectors were mutagenized in XL1-red (Agilent) as per manufacturer’s instructions. XL1-red transformants were washed off the transformation plate, grown overnight in 50 ml LB with ampicillin and the resulting mutant plasmid pool was purified by midiprep (Promega). The mutant plasmid pool was transformed to BY4742 using the high-efficiency LiOAC/PEG method (Gietz and Schiestl 2007). Four transformation reactions were performed per screen, each using 1 μg of plasmid pool per reaction. One reaction was split in three parts and plated onto different media types to quantify the transformable plasmids (SD- uracil + 2% glucose), loss-of-function mutants (SD- uracil +2% galactose), and missense loss-of-function mutants (SD-uracil/histidine + 2% galactose) and incubated for 2-4 days at 30°C. The remaining transformation reactions were plated onto 150 mm SD-uracil/histidine + 2% galactose plates and allowed to grow until colonies appeared (3-4 days). Plasmids were rescued from missense loss-of-function mutants and transformed to the *E. coli* Top10 strain before sequencing using primers in the vector and effector, if required (Table S1). Details of the sequenced mutants are shown in Table S2.

### Analysis of mutant fitness

Liquid growth assays were used to assess the effect of missense loss-of-function mutations on yeast fitness as described (Urbanus *et al*. 2016) with the following modifications. Overnight cultures of freshly transformed BY4742 with empty vector control, pYES2 NT/A effector-HIS3 wild-type and mutants were diluted 100-fold into 100 μl of SD-ura/2% galactose and grown with Breathe-Easy adhesive seals (EK scientific) in a CellGrower robot (S&P robotics) at 30°C with intermittent shaking. Yeast growth was monitored for 30 h by measuring the OD_620_ every 15 min. Growth fitness was calculated as the ratio of the area under the curve (AUC) of a effector-expressing strain over an empty vector control after 30 h using the R package GrowthCurver (Sprouffske and Wagner 2016). The average AUC ratio and standard deviation was calculated from three technical replicates.

### HHpred analysis and sequence alignments

The amino acid sequence of RavK and SdbA (amino acid residues 528-1116) were submitted to the HHpred server (https://toolkit.tuebingen.mpg.de/#/) (Zimmermann *et al*. 2018) analyzed using MSA generation HHblits Uniclust20_2017_07 and Uniprot20_2016_02, respectively and otherwise default parameters. The resulting alignments were visualized using Boxshade (https://embnet.vital-it.ch/software/BOX_form.html). Amino acid sequence alignments of SdbB with its orthologs or with SidB were generated using T-coffee (Tommaso *et al*. 2011) and visualized using Jalview (Waterhouse *et al*. 2009) or Boxshade.

### Nucleotide sugar donor specificity of the SdbA C-terminal domain

The gene fragment corresponding to SdbA (Lpg0275) residues 510-1050 was PCR amplified from *Legionella pneumophila* str Philadelphia-1 genomic DNA and inserted into the pMCSG53 plasmid (Eschenfeldt *et al*. 2013) by ligation independent cloning, providing an N-terminal 6xHIS-TEV tag. The point mutant E963A was prepared by site-directed mutagenesis using QuikChange™ site-directed mutagenesis kit (Stratagene) according to the manufacturer’s protocol. Plasmids were sequenced and transformed to the *E. coli* BL21 DE Gold strain for purification. Recombinant proteins were purified to near homogeneity (>95%) using Ni-chelate affinity chromatography on Ni-NTA Superflow resin (Qiagen) using standard protocols. Cultures were grown in TB and expression was induced at an OD_595_ of 0.8 with 0.4 mM IPTG overnight at 16 °C. Cells were harvested by centrifugation at 9,300× *g*, resuspended in 50 mM HEPES pH 7.5, 400 mM NaCl, 5% glycerol, 5 mM imidazole, and lysed by sonication. Lysates were clarified by centrifugation at 21,000× *g* at 4°C and loaded onto gravity flow Ni-NTA agarose columns (Qiagen), followed by washing with 50 mM HEPES pH 7.5, 400 mM NaCl, 5% glycerol, 30 mM imidazole. Proteins were eluted using 50 mM HEPES pH 7.5, 400 mM NaCl, 5% glycerol, 250 mM imidazole and flash-frozen in liquid nitrogen for storage at −80°C. The purity of the protein samples was assessed by SDS-PAGE and visualized by Coomassie Brilliant Blue R.

The nucleotide sugar donor specificity of SdbA 510-1050 was assayed using the UDP-Glo Glycosyltransferase Assay (Promega) according to manufacturer’s protocol. Briefly, 0.09 µM of purified wild-type and E963A mutant SbdA 510-1050 protein was incubated with 100 µM UDP-glucose, UDP-glcNAc, UDP-glucuron, UDP-galactose or UDP-galNAc for 1h at 30°C in 50 mM HEPES, pH 7.5, 100 mM KCl, 2 mM MgCl_2_, 1 mM MnCl_2._ The hydrolase activity of the UDP-substrate was detected as the release of UDP by the UDP-Glo assay (Promega) after 20 min of incubation with UDP-Glo detection reagent. Luminescence was measured using a SpectraMax M2 plate reader. Three technical repeats were performed per reaction.

The V_max_, K_m_ and k_cat_ for wild-type SdbA 510-1050 with UDP-glcNAc was determined by incubating 0.16 µM SdbA 510-1050 with UDP-glcNAc concentration range of 0.0039 – 2 mM for 1h at 30°C in 50 mM HEPES, pH 7.5, 100 mM KCl, 2 mM MgCl_2_, 1 mM MnCl_2._ Three technical repeats were performed per reaction. Kinetic parameters were determined by non-linear curve fitting from the Michaelis Menten plot using GraphPad Prism (version 5.00 for Windows, GraphPad Software).

## Acknowledgments

We thank Beth Nicholson, Morgan Petersen and John McPherson, for their suggestions and careful reading of the manuscript and Kamran Rizzolo for help with cloning. This work was supported by a Project Grant (AWE) from the Canadian Institutes of Health Research (PJT- 162256).

---

**Figure S1:**
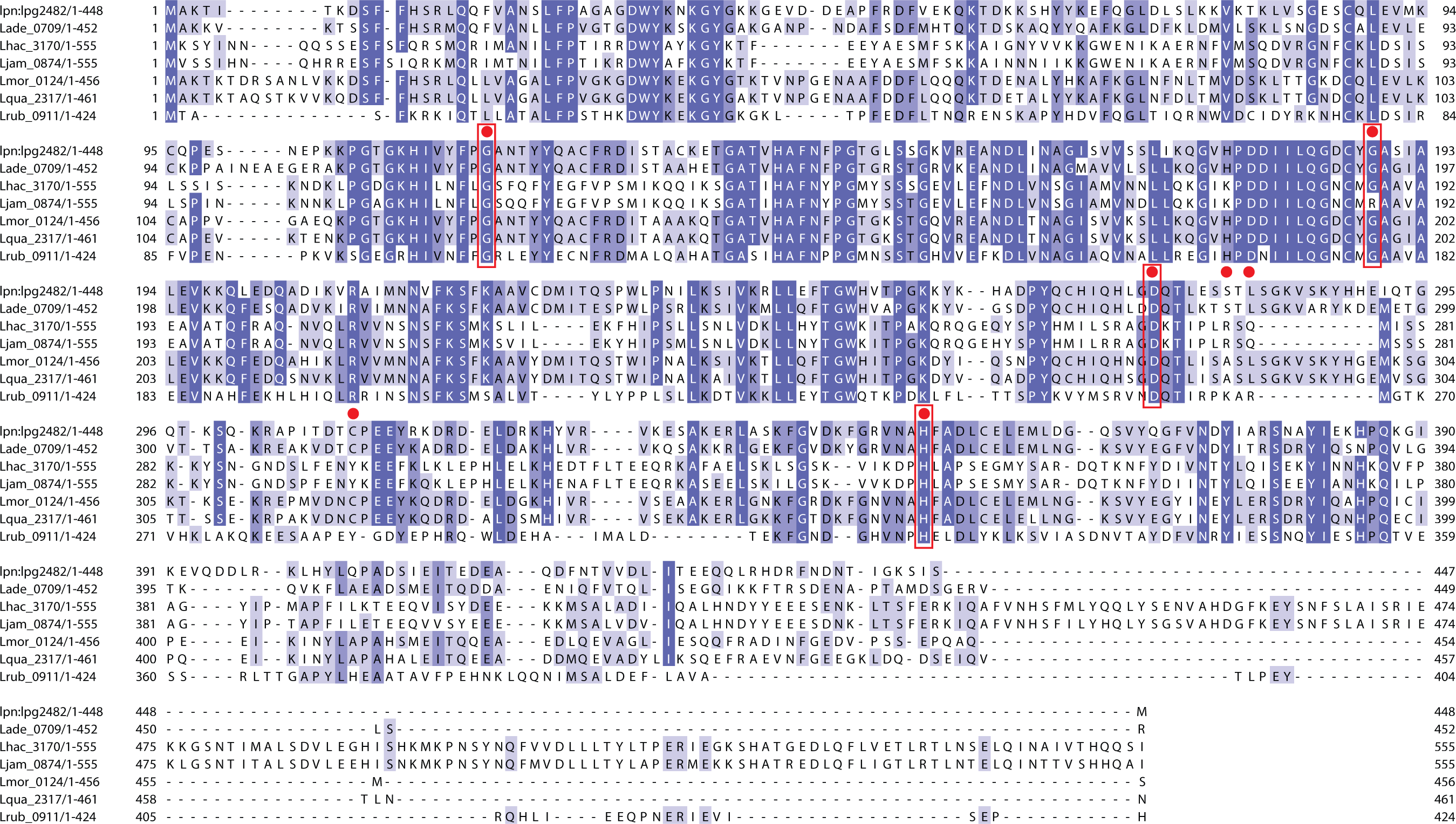
Random mutagenesis targets conserved residues in SdbB orthologs from seven *Legionella* species. The sequences of SdbB orthologs identified by Burstein et al (Nat Genet 48(2): 167-175, 2016) were aligned using T-coffee, visualized with Jalview and coloured by % identity. The missense mutations identified by the random mutagenesis screen are indicated with a red closed circle above the SdbB (lpg2482) sequence. Four of the seven mutations target invariant residues; G116, G189, D273 and H351 shown in red boxes.

**Figure S2:**
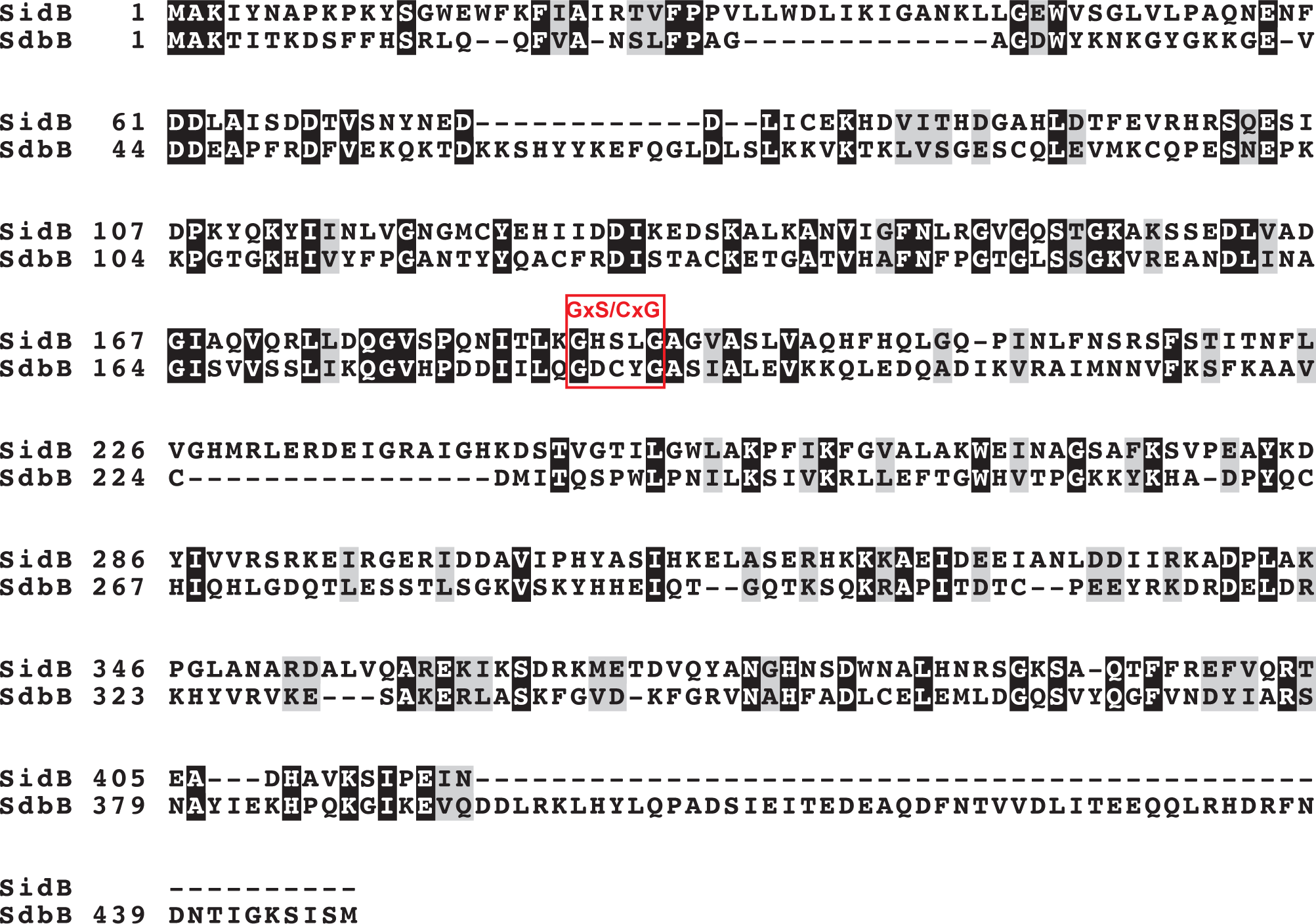
**SidB and SdbB amino acid sequence alignment**. The SidB (lpg1642) and SdbB (lpg2482) were aligned with T-coffee and visualized with Boxshade, where identical residues are shown in black and similar residues in grey. The active site motif GxS/CxG predicted by NCBI conserved domain search (Marchler-Bauer A et al.,2017, Nucleic Acids Res.45(D)200-3.) for SidB is indicated, suggesting that SdbB C187 is part of the active site catalytic triad.

**Figure S3.**
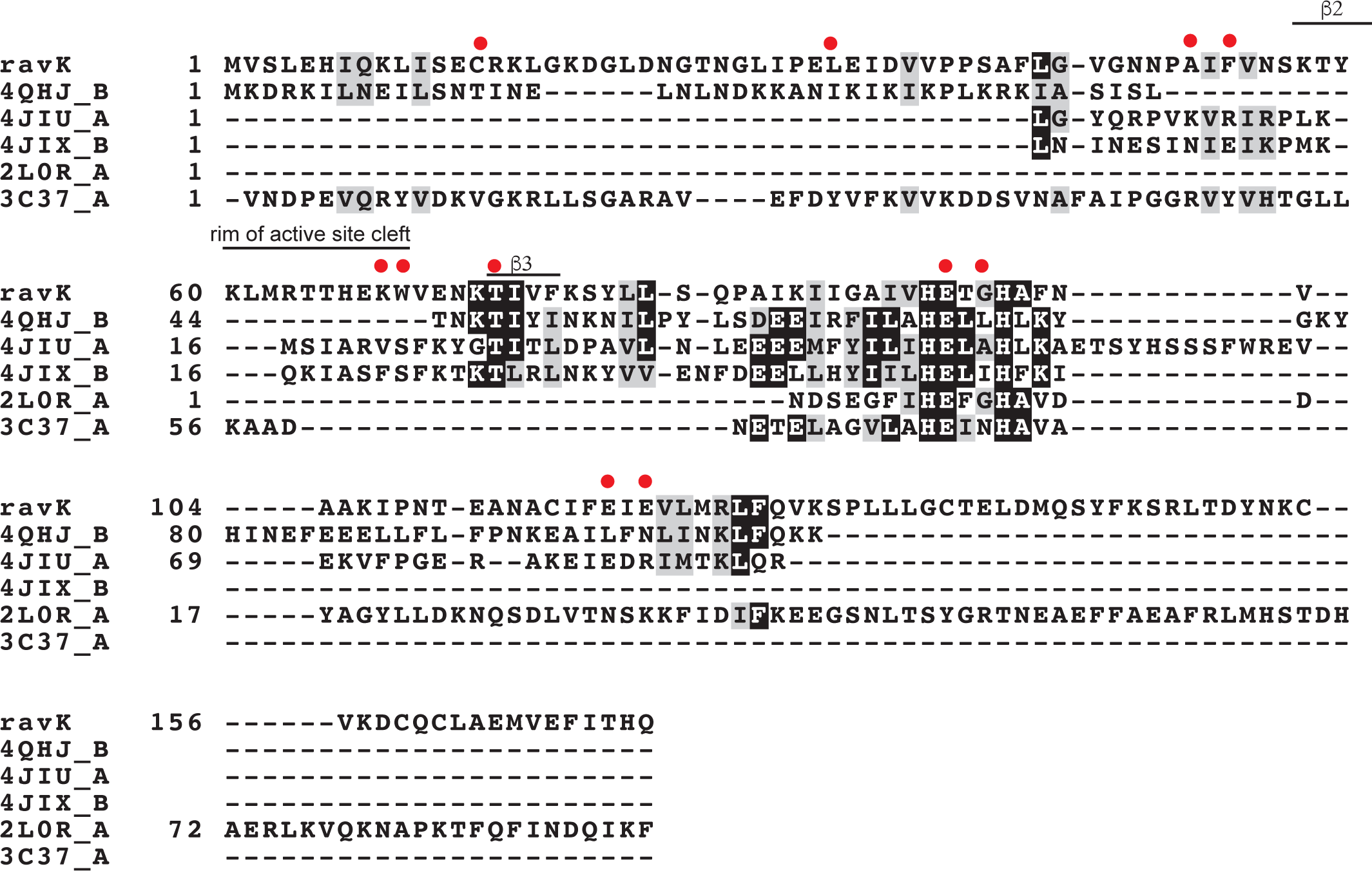
HHpred alignment of RavK with metalloproteases. Alignment of the HHpred top hits; MJ1213 (4QHJ), proabylysin (4JIU), projannalysin (4JIX) anthrax lethal factor (2L0R) and a zinc-dependent endopeptidase of the M84 family (3C37) with RavK is shown where grey and black background indicate the level of amino acid residue conservation as 50% or more similarity or 50% or more identity, respectively. Structural elements of proabylysin (4JIU) are indicated on top of the alignment and mutations that abolish RavK activity are indicated by a red closed circle.

**Table S1:**
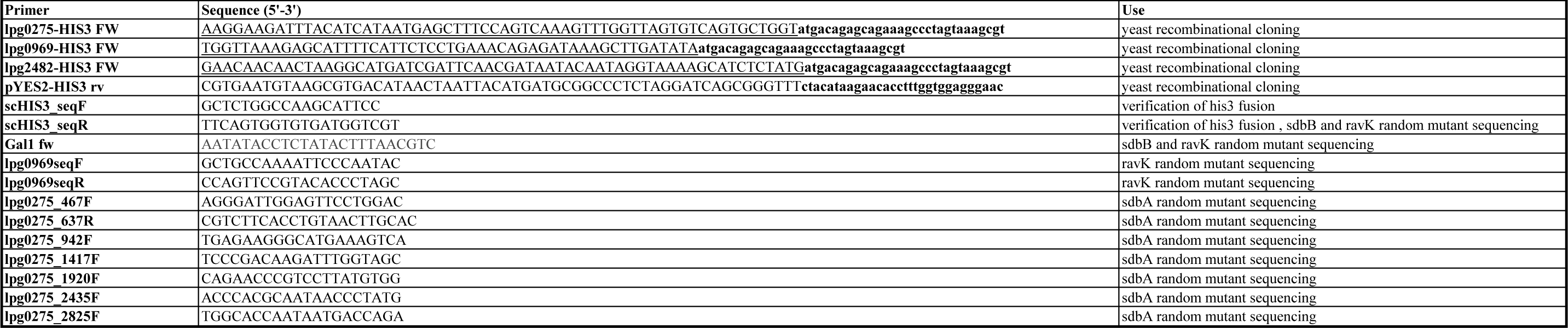
Primers used for yeast recombinational cloning and sequencing. In the yeast recombinational cloning primers the effector sequence is underlined, the HIS3 sequence in bold and the pYES2 vector sequence unchanged.

**Table S2.** Random mutagenesis mutants for RavK, SdbA and SdbB. Mutants suppressing the growth defect of RavK,SdbA and SdbB in yeast were selected on SD-gal-ura (induced expression and plasmid selection) or SD-gal-ura-his (induced expression, plasmid selection and full-length fusion protein selection respectively). This table summarizes the identified mutations, the type of mutation and the consequences of the mutation.

| Mutant no | Locus no | Gene name | Selection | Nucleotide mutation | Type of mutation | Amino acid mutation(s) | 1st mutation | 2nd mutation |
| --- | --- | --- | --- | --- | --- | --- | --- | --- |
| 1 | lpg0275 | sdbA | + gal, -ura-his | 367G>A, 1624G>A | amino acid substitution | V123M, G542R | 123 | 542 |
| 2 | lpg0275 | sdbA | + gal, -ura-his | 1622G>A, 452C>T | amino acid substitution | S151L, G541E | 151 | 541 |
| 3 | lpg0275 | sdbA | + gal, -ura-his | 1222G>A, 1622G>A | amino acid substitution | V408I, G541E | 408 | 541 |
| 4 | lpg0275 | sdbA | + gal, -ura-his | 1622G>A | amino acid substitution | G541E | 541 |  |
| 5 | lpg0275 | sdbA | + gal, -ura-his | 1621G>A | amino acid substitution | G541E | 541 |  |
| 6 | lpg0275 | sdbA | + gal, -ura-his | 1622G>A | amino acid substitution | G541E | 541 |  |
| 7 | lpg0275 | sdbA | + gal, -ura-his | 1622G>A | amino acid substitution | G541E | 541 |  |
| 8 | lpg0275 | sdbA | + gal, -ura-his | 1622G>A | amino acid substitution | G541E | 541 |  |
| 9 | lpg0275 | sdbA | + gal, -ura-his | 1625G>A | amino acid substitution | G542E | 542 |  |
| 10 | lpg0275 | sdbA | + gal, -ura-his | 1630G>A, 2378C>T | amino acid substitution | G544R, P793L | 544 | 793 |
| 11 | lpg0275 | sdbA | + gal, -ura-his | 1630G>A, 2826G>A | amino acid substitution | G544R, M942I | 544 | 942 |
| 12 | lpg0275 | sdbA | + gal, -ura-his | 1634A>G | amino acid substitution | H545R | 545 |  |
| 13 | lpg0275 | sdbA | + gal, -ura-his | 1633C>T | amino acid substitution | H545Y | 545 |  |
| 14 | lpg0275 | sdbA | + gal, -ura-his | 1639T>C | amino acid substitution | S547P | 547 |  |
| 15 | lpg0275 | sdbA | + gal, -ura-his | 2275G>A | amino acid substitution | D759N | 759 |  |
| 16 | lpg0275 | sdbA | + gal, -ura-his | 2275G>A | amino acid substitution | D759N | 759 |  |
| 17 | lpg0275 | sdbA | + gal, -ura-his | 2731C>A, 2866G>A | amino acid substitution | G911R, G956R | 911 | 956 |
| 18 | lpg0275 | sdbA | + gal, -ura-his | 2870G>A | amino acid substitution | G957D | 957 |  |
| 19 | lpg0275 | sdbA | + gal, -ura-his | 2873G>A | amino acid substitution | G958D | 958 |  |
| 20 | lpg0275 | sdbA | + gal, -ura-his | 2887G>A | amino acid substitution | E963K | 963 |  |
| 21 | lpg0275 | sdbA | + gal, -ura-his | 2887G>A | amino acid substitution | E963K | 963 |  |
| 22 | lpg0275 | sdbA | + gal, -ura-his | 2887G>A | amino acid substitution | E963K | 963 |  |
| 23 | lpg0275 | sdbA | + gal, -ura-his | 2980G>A | amino acid substitution | E994K | 994 |  |
| 24 | lpg0275 | sdbA | + gal, -ura-his | 2980G>A | amino acid substitution | E994K | 994 |  |
| 25 | lpg0969 | ravK | + gal, -ura-his | 41G>A | amino acid substitution | C14Y | 14 |  |
| 26 | lpg0969 | ravK | + gal, -ura-his | 98T>C | amino acid substitution | L33S | 33 |  |
| 27 | lpg0969 | ravK | + gal, -ura-his | 152C>T | amino acid substitution | A51V | 51 |  |
| 28 | lpg0969 | ravK | + gal, -ura-his | 158T>C | amino acid substitution | F53S | 53 |  |
| 29 | lpg0969 | ravK | + gal, -ura-his | 203A>C | amino acid substitution | K68T | 68 |  |
| 30 | lpg0969 | ravK | + gal, -ura-his | 205T>A | amino acid substitution | W69R | 69 |  |
| 31 | lpg0969 | ravK | + gal, -ura-his | 220A>G | amino acid substitution | T74A | 74 |  |
| 32 | lpg0969 | ravK | + gal, -ura-his | 286G>A | amino acid substitution | E96K | 96 |  |
| 33 | lpg0969 | ravK | + gal, -ura-his | 286G>A | amino acid substitution | E96K | 96 |  |
| 34 | lpg0969 | ravK | + gal, -ura-his | 287A>G | amino acid substitution | E96G | 96 |  |
| 35 | lpg0969 | ravK | + gal, -ura-his | 293G>T | amino acid substitution | G98V | 98 |  |
| 36 | lpg0969 | ravK | + gal, -ura-his | 331G>A | amino acid substitution | E111K | 111 |  |
| 37 | lpg0969 | ravK | + gal, -ura-his | 354A>C | amino acid substitution | E118D | 118 |  |
| 38 | lpg2482 | sdbB | +gal,-ura | 33_34insT | frameshift | H12SfsX1 | 12 |  |
| 39 | lpg2482 | sdbB | +gal,-ura | 33_34insT | frameshift | H12SfsX1 | 12 |  |
| 40 | lpg2482 | sdbB | +gal,-ura | 33_34insT | frameshift | H12SfsX1 | 12 |  |
| 41 | lpg2482 | sdbB | +gal,-ura | 119delA | frameshift | K40RfsX33 | 39 |  |
| 42 | lpg2482 | sdbB | +gal,-ura | 119delA | frameshift | K40RfsX33 | 39 |  |
| 43 | lpg2482 | sdbB | +gal,-ura | 166_167insA | frameshift | Q56TfsX4 | 56 |  |
| 44 | lpg2482 | sdbB | +gal,-ura | 172_173insA | frameshift | T58NfsX2 | 58 |  |
| 45 | lpg2482 | sdbB | +gal,-ura | 172_173insA | frameshift | T58NfsX2 | 58 |  |
| 46 | lpg2482 | sdbB | +gal,-ura | 172_173insA | frameshift | T58NfsX2 | 58 |  |
| 47 | lpg2482 | sdbB | +gal,-ura | 172delA | frameshift | T58PfsX15 | 58 |  |
| 48 | lpg2482 | sdbB | +gal,-ura | 265C>T | premature stop codon | Q89AMBER | 89 |  |
| 49 | lpg2482 | sdbB | +gal,-ura | 312delA | frameshift | K104NfsX56 | 104 |  |
| 50 | lpg2482 | sdbB | +gal,-ura | 407G>A, 469_470insG | amino acid substitution and frameshift | G136E, G157AfsX2 | 136 | 157 |
| 51 | lpg2482 | sdbB | +gal,-ura | 555_556insG | frameshift | G185=fsX1 | 185 |  |
| 52 | lpg2482 | sdbB | +gal,-ura | 594_595insA | frameshift | Q199AfsX21 | 199 |  |
| 53 | lpg2482 | sdbB | +gal,-ura | 818A>G | amino acid substitution | D273G | 273 |  |
| 54 | lpg2482 | sdbB | +gal,-ura | 905_906insA | frameshift | R302KfsX17 | 302 |  |
| 55 | lpg2482 | sdbB | +gal,-ura | 905_906insA | frameshift | R302KfsX17 | 302 |  |
| 56 | lpg2482 | sdbB | +gal,-ura | 905delA | frameshift | R302GfsX40 | 302 |  |
| 57 | lpg2482 | sdbB | +gal,-ura | 1094C>A | premature stop codon | S365AMBER | 365 |  |
| 58 | lpg2482 | sdbB | + gal, -ura-his | 347G>A | amino acid substitution | G116E | 116 |  |
| 59 | lpg2482 | sdbB | + gal, -ura-his | 347G>A | amino acid substitution | G116E | 116 |  |
| 60 | lpg2482 | sdbB | + gal, -ura-his | 346G>A | amino acid substitution | G116R | 116 |  |
| 61 | lpg2482 | sdbB | + gal, -ura-his | 347G>A | amino acid substitution | G116E | 116 |  |
| 62 | lpg2482 | sdbB | + gal, -ura-his | 347G>A | amino acid substitution | G116E | 116 |  |
| 63 | lpg2482 | sdbB | + gal, -ura-his | 347G>A | amino acid substitution | G116E | 116 |  |
| 64 | lpg2482 | sdbB | + gal, -ura-his | 347G>A | amino acid substitution | G116E | 116 |  |
| 65 | lpg2482 | sdbB | + gal, -ura-his | 347G>A | amino acid substitution | G116E | 116 |  |
| 66 | lpg2482 | sdbB | + gal, -ura-his | 566G>A | amino acid substitution | G189E | 189 |  |
| 67 | lpg2482 | sdbB | + gal, -ura-his | 566G>A | amino acid substitution | G189E | 189 |  |
| 68 | lpg2482 | sdbB | + gal, -ura-his | 836C>T | amino acid substitution | S279L | 279 |  |
| 69 | lpg2482 | sdbB | + gal, -ura-his | 836C>T | amino acid substitution | S279L | 279 |  |
| 70 | lpg2482 | sdbB | + gal, -ura-his | 836C>T | amino acid substitution | S279L | 279 |  |
| 71 | lpg2482 | sdbB | + gal, -ura-his | 842T>C | amino acid substitution | L281S | 281 |  |
| 72 | lpg2482 | sdbB | + gal, -ura-his | 842T>C | amino acid substitution | L281S | 281 |  |
| 73 | lpg2482 | sdbB | + gal, -ura-his | 842T>C | amino acid substitution | L281S | 281 |  |
| 74 | lpg2482 | sdbB | + gal, -ura-his | 926G>A | amino acid substitution | C309Y | 309 |  |
| 75 | lpg2482 | sdbB | + gal, -ura-his | 926G>A | amino acid substitution | C309Y | 309 |  |
| 76 | lpg2482 | sdbB | + gal, -ura-his | 926G>A | amino acid substitution | C309Y | 309 |  |
| 77 | lpg2482 | sdbB | + gal, -ura-his | 1051C>T | amino acid substitution | H351Y | 351 |  |

